# Improved *De Novo* Peptide Binder Design with Target-Conditioned Inverse Folding

**DOI:** 10.1101/2025.10.28.685072

**Authors:** Elliot Layne, Ashlin K. Kanawaty, Aron Broom, Alexander Kitaygorodsky, Anita K. Nivedha, Parth Vora, Caroline Woffindale, Sarah Hailstone, Eirini Sachouli, Ella Halcrow, Gillian Donachie, Michelle Adsett, Glenn L. Butterfoss, Mark Fingerhuth

## Abstract

Inverse protein folding methods have become central to the computational design of *de novo* proteins, but existing models struggle when tasked with generating high-affinity peptide binders. By combining peptide-specific finetuning with a novel decoding order strategy, we enhance pocket conditioning and enable more accurate sequence design for peptide-binding interfaces. Our approach delivers gains in computational metrics, increasing sequence recovery and improving *in silico* binder design success rate by 16% *™* 30%. *In vitro* validation finds that our method greatly improves the success rate of designing novel peptide agonists of the OPRM1 receptor, generating at least twice as many top-ranking agonists as the prevailing standard method ProteinMPNN.

## 1 Introduction

Generative protein design methods play an increasingly significant role in computational drug discovery efforts. Binder design methods such as RFDiffusion [Watson et al., 2023] and BindCraft [Pacesa et al., 2025] have a growing track record of experimentally validated success. These methods exhibit a characteristic two-step generative process, in which the first step is to generate a backbone conformation, conditioned on a target receptor structure. In the second step, this backbone is passed as input to the inverse folding model ProteinMPNN [Dauparas et al., 2022], which predicts a sequence of amino acids that will adopt the specified backbone conformation within the target pocket.

An influential message-passing graph neural network (MPNN) algorithm for protein design was introduced by Ingraham et al. [2019], in which the protein backbone is represented as a graph and residue-level information is propagated through edges to capture local geometric context. This work demonstrated that GNN-based generative models could learn structure–sequence relationships and produce plausible protein sequences conditioned on a fixed backbone. Building on this foundation, Dauparas et al. [2022] refined the message passing framework by incorporating richer geometric edge features and introducing a random decoding order during sequence generation. Their method, ProteinMPNN, significantly advanced the field, achieving higher sequence recovery rates and more stable designs compared to prior approaches. It remains a benchmark model for backbone-conditioned protein sequence design.

Peptides are a less common, but important, class of therapeutics that possess properties at the intersection of small molecules and proteins. Peptides have higher specificity than, and generally similar affinity to, small molecules [Giacomo Rossino, 2023], better tissue penetration and internalization than antibodies [Sri Murugan Poongkavithai Vadevoo, 2023], and lower immunogenicity and toxicity compared to other biologics [Lamers, 2022]. While there is a wealth of protein structure data, peptide structural data is far less common. Additionally, peptides tend to be more flexible than proteins. The scarcity of data, combined with the conformational heterogeneity of many peptides, makes structure-based design of peptides challenging for methods developed for proteins, including ProteinMPNN.

To address this need, we sought to finetune ProteinMPNN for the task of generating peptide sequences conditionally on a target binding site, referring to our updated model as PeptideMPNN. We present both computational and *in vitro* results demonstrating that both pretraining and finetuning with elevated backbone noise improves peptide binder sequence design.

Further, we present a novel decoding order for autoregressive inverse folding models, optimized for binder design by prioritizing residues located at the binding interface. We find that this contact decoding order improves *in silico* performance of *de novo* binder design methods in comparison to the commonly used random order decoding.

There have been several attempts to re-purpose ProteinMPNN for specific applications such as designing HLA peptide binders [Xu et al., 2024], antibodies [Sun et al., 2025, Dreyer et al., 2023, Shanehsazzadeh], and small-molecule binding proteins [Dauparas et al., 2025]. While the aforementioned methods collectively highlight the expanding role of graph– and deep-learning methods in tailoring protein models for therapeutic applications, there remains a need for improved methods for general peptide binder sequence design from structure.

## 2 Methods

### Data preparation

We constructed a dataset from the union of all peptide-protein complexes included within the Peptide Binding DataBase (PepBDB) [Wen et al., 2019], the datasets used by Wang et al. [2024] and Kong et al. [2024], and any additional peptide-protein complexes deposited to the Protein Data Bank (PDB) before January 2025 [Berman et al., 2000]. All examples in which the peptide chain was of length less than 5 or greater than 30 residues were removed from consideration. PDB structures were restricted to samples solved by X-ray crystallography or Cryo-EM with a minimum resolution of 3.5 angstroms.

The resulting dataset contained 11,898 peptide-protein complexes. The peptide sequences were clustered with MMseqs2 [Steinegger and Söding, 2017], ensuring a maximum 40% sequence similarity between any pair of peptides across clusters. We randomly selected a subset of clusters containing a total of 1000 peptides as a held out test set. The remaining clusters were used as training data.

### Finetuning regime

Given that peptides have more highly varying conformations than proteins, we experimented with independently varying the degree of backbone noise added to training samples during both pretraining and finetuning. To optimize the degree of backbone noise, we finetuned each of the four base ProteinMPNN models, pretrained with Gaussian noise variances *ϵ∈* 0.02, 0.1, 0.2, 0.3. Each base model was further finetuned on our peptide complex dataset using the same set of *ϵ* values.

For every base–finetuning combination, we performed 10 replicates and report averages across runs. For each finetuning replicate, the training clusters were randomly split into a train dataset (90%) and a validation set (10%).

All models were finetuned for 200 epochs, using the AdamW optimizer [Loshchilov and Hutter, 2017] with an initial learning rate of 5*×*10^*™*5^. We maintained the dropout and label smoothing rates of 10% during training utilized by Dauparas et al. [2022] and Ingraham et al. [2019].

Early experimentation explored finetuning utilizing the “NoamOpt” learning rate schedule described by Vaswani et al. [2017], as used by Dauparas et al. [2022] during training of base models. These results are further described in Appendix A.3. We note the *in vitro* experimentation described in Section 3.1 was performed using this earlier finetuning regime.

### Contact-based decoding order

Dauparas et al. [2022] showed that a randomized decoding order of fixed residues followed by masked residues had slightly better sequence recovery than simple left-to-right decoding. In our context of designing a peptide binding sequence for a fixed target, and given permutation function, *π*, we can express the random ProteinMPNN decoding order function, *D*_*rand*_, as

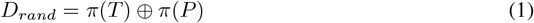

Where *T* represents fixed residues in the visible chain (the target), and *P* represents designable residues in the masked chain (the peptide). We note that the decoding order of the fixed residues is relevant as it affects the message-passing algorithm.

We hypothesized that designable, contacting residues involved in binding are more constrained in their amino acid identity than designable, non-contacting residues due to their interaction with a target. We therefore attempted to further improve sequence recovery by decoding masked residues in close contact with the target before masked residues that do not make contact.

Using the backbone structure, we were able to compute pairwise distances between C-*α* atoms from different chains. We defined contacts as any pair of residues with a distance between C-*α* atoms below the threshold, *d*_*c*_ = 8 angstroms.

We were then able to classify residues into the following groups, with the groups being decoded in order:

1. Fixed. In our case, these are exclusively in the target, (*T*)
2. Masked and contacting in the peptide, *P*_*C*_
3. Masked and non-contacting in the peptide, *P*_*NC*_

Figure 4 in Appendix A.2 illustrates the labeling of these groups on an example structure. As with ProteinMPNN’s randomized decoding implementation, the order of residues within each group was randomized to create a contact-based decoding order, *D*_*cont*_.

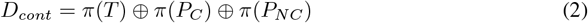

## 3 Experimentation

### Sequence recovery and perplexity across finetuning regimes

We evaluated the performance of the base and finetuned models with regards to sequence recovery and perplexity on our held out test set (Table 1 and Table 2).

**Table 1:**
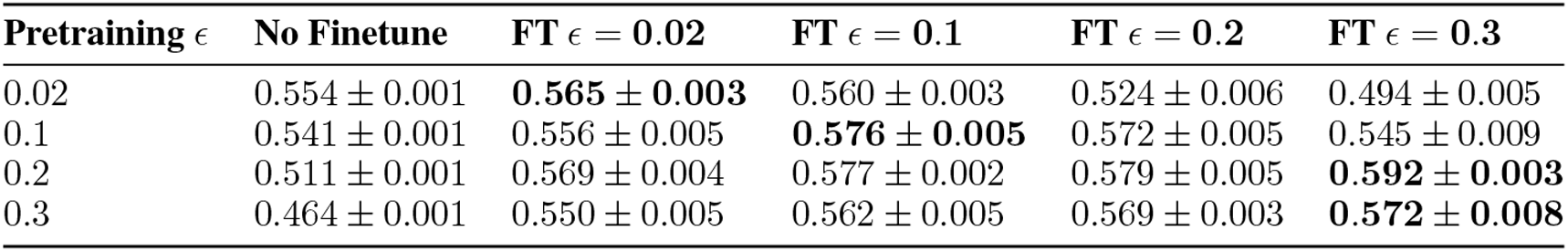
Peptide sequence recovery across pretraining and finetuning *ϵ* values.

**Table 2:** Peptide sequence perplexity across pretraining and finetuning *ϵ* values.

| Pretraining $\epsilon$ | No Finetune | FT $\epsilon = 0.02$ | FT $\epsilon = 0.1$ | FT $\epsilon = 0.2$ | FT $\epsilon = 0.3$ |
| --- | --- | --- | --- | --- | --- |
| 0.02 | **4.710 ± 0.004** | 5.836 ± 0.075 | 4.989 ± 0.074 | 5.561 ± 0.095 | 6.048 ± 0.072 |
| 0.1 | 4.940 ± 0.003 | 6.022 ± 0.111 | 5.196 ± 0.148 | **4.903 ± 0.096** | 4.955 ± 0.099 |
| 0.2 | 5.362 ± 0.003 | 5.819 ± 0.126 | 5.448 ± 0.107 | 4.919 ± 0.088 | **4.698 ± 0.046** |
| 0.3 | 5.886 ± 0.005 | 5.970 ± 0.153 | 5.340 ± 0.057 | 4.916 ± 0.064 | **4.671 ± 0.071** |

Finetuning improved sequence recovery across all base models. The base model pretrained with low-noise setting *ϵ* = 0.02 exhibited improved sequence recovery when finetuned with a matching value for *ϵ*. All other base models exhibited improved sequence recovery for all choices of *ϵ* during finetuning. Globally, the highest sequence recovery was exhibited by models that were both pretrained and finetuned with high levels of backbone noise.

Finetuning did not improve sequence perplexity for the base model pretrained with low backbone noise (*ϵ≤* 0.1). All other base models exhibited improved perplexity after finetuning, with the best sequence perplexity results exhibited by models that were both pretrained and finetuned with high levels of backbone noise.

Across both metrics, pretraining with *ϵ ≥* 0.2 and finetuning with *ϵ* = 0.3 exhibited optimal results. These findings suggests that training inverse folding models with elevated backbone noise can improve generalization performance on peptides, possibly due to increasing robustness to conformational variability.

### Decoding order effects on *de novo* binder design

We evaluated the ability of a contact-based decoding order to improve the design of *de novo* peptide sequences conditioned on a target binding site, using OPRM1 as a case study. We diffused 100 peptide backbones of length 16-20 residues using RFDiffusion [Watson et al., 2023], conditioned to bind four expert-selected hotspots within the binding pocket. We note that in order to assess the effects of contact decoding, the length range was extended beyond what was explored in Section 3.1. This ensured that a portion of the peptide backbone would be placed outside of the binding pocket, guaranteeing the existence of contacting (ie. in the binding pocket) and non-contacting (ie. outside the binding pocket) peptide residues.

We retained 40 diffused backbones that made contact with the binding pocket. For each backbone, we generated 50 peptide sequences with PeptideMPNN using both random and contact-first decoding orders with a sampling temperature of 0.2.

We refolded each *de novo* peptide sequence together with the target using AlphaFold2 (AF2), using the “initial guess” protocol as described by Bennett et al. [2023]. We compared the number of *in silico* “hits”, defined by the refolded complex having an AF2 iPAE score below 10 or 15 (Figure 1). The use of contact decoding increased hit rate by 16% and 30% respectively, suggesting there is a substantial benefit to decoding residues in contact with the binding site first during *de novo* design. Structure and sequence analysis of the *in silico* hits can be found in A.2.3, Figures 6 and 7.

**Figure 1.**
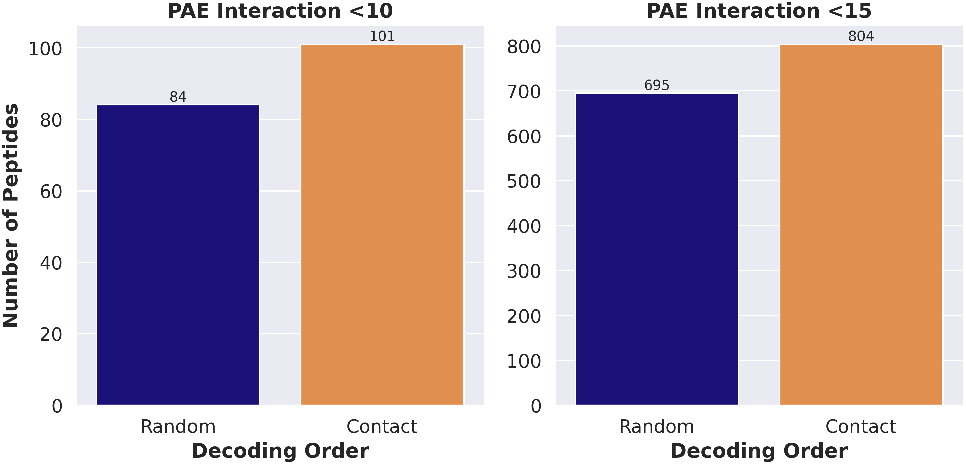
The number of *de novo* sequences generated by each decoding order strategy which resulted in a refolded complex with AlphaFold2 iPAE below 10 (left panel) or 15 (right panel). The contact decoding order results in an increased number of hits at both thresholds.

In addition to improving *de novo* binder design, we assessed the impact of contact decoding on ground-truth sequence recovery, finding that it exhibited no loss in performance in comparison with random order decoding (see Appendix A.2.2, Tables 3 and 4).

**Table 3:** Peptide sequence recovery across pretraining and finetuning *ϵ* values.

| Pretraining $\epsilon$ | No Finetune | FT $\epsilon = 0.02$ | FT $\epsilon = 0.1$ | FT $\epsilon = 0.2$ | FT $\epsilon = 0.3$ |
| --- | --- | --- | --- | --- | --- |
| **Random decoding order** |  |  |  |  |  |
| 0.02 | $0.554 \pm 0.001$ | **$0.565 \pm 0.003$** | $0.560 \pm 0.003$ | $0.524 \pm 0.006$ | $0.494 \pm 0.005$ |
| 0.1 | $0.541 \pm 0.001$ | $0.556 \pm 0.005$ | **$0.576 \pm 0.005$** | $0.572 \pm 0.005$ | $0.545 \pm 0.009$ |
| 0.2 | $0.511 \pm 0.001$ | $0.569 \pm 0.004$ | $0.577 \pm 0.002$ | $0.579 \pm 0.005$ | **$0.592 \pm 0.003$** |
| 0.3 | $0.464 \pm 0.001$ | $0.550 \pm 0.005$ | $0.562 \pm 0.005$ | $0.569 \pm 0.003$ | **$0.572 \pm 0.008$** |
| **Contact decoding order** |  |  |  |  |  |
| 0.02 | $0.551 \pm 0.001$ | **$0.566 \pm 0.003$** | $0.563 \pm 0.003$ | $0.519 \pm 0.007$ | $0.493 \pm 0.006$ |
| 0.1 | $0.541 \pm 0.001$ | $0.558 \pm 0.005$ | **$0.578 \pm 0.004$** | $0.575 \pm 0.006$ | $0.545 \pm 0.009$ |
| 0.2 | $0.509 \pm 0.001$ | $0.570 \pm 0.005$ | $0.576 \pm 0.003$ | $0.579 \pm 0.003$ | **$0.591 \pm 0.003$** |
| 0.3 | $0.464 \pm 0.002$ | $0.548 \pm 0.006$ | $0.562 \pm 0.006$ | **$0.570 \pm 0.005$** | $0.570 \pm 0.008$ |

**Table 4:** Peptide sequence perplexity across pretraining and finetuning *ϵ* values.

| Pretraining $\epsilon$ | No Finetune | FT $\epsilon = 0.02$ | FT $\epsilon = 0.1$ | FT $\epsilon = 0.2$ | FT $\epsilon = 0.3$ |
| --- | --- | --- | --- | --- | --- |
| **Random decoding order** |  |  |  |  |  |
| 0.02 | **4.710 ± 0.004** | 5.836 ± 0.075 | 4.989 ± 0.074 | 5.561 ± 0.095 | 6.048 ± 0.072 |
| 0.1 | 4.940 ± 0.003 | 6.022 ± 0.111 | 5.196 ± 0.148 | **4.903 ± 0.096** | 4.955 ± 0.099 |
| 0.2 | 5.362 ± 0.003 | 5.819 ± 0.126 | 5.448 ± 0.107 | 4.919 ± 0.088 | **4.698 ± 0.046** |
| 0.3 | 5.886 ± 0.005 | 5.970 ± 0.153 | 5.340 ± 0.057 | 4.916 ± 0.064 | **4.671 ± 0.071** |
| **Contact decoding order** |  |  |  |  |  |
| 0.02 | **4.705 ± 0.004** | 5.835 ± 0.065 | 4.981 ± 0.059 | 5.612 ± 0.103 | 6.072 ± 0.089 |
| 0.1 | 4.941 ± 0.007 | 5.980 ± 0.108 | 5.152 ± 0.142 | **4.881 ± 0.100** | 4.968 ± 0.087 |
| 0.2 | 5.351 ± 0.005 | 5.819 ± 0.137 | 5.445 ± 0.091 | 4.921 ± 0.075 | **4.718 ± 0.049** |
| 0.3 | 5.873 ± 0.009 | 5.987 ± 0.158 | 5.353 ± 0.067 | 4.902 ± 0.075 | **4.692 ± 0.065** |

### 3.1 *In vitro* validation

We undertook a large-scale binder design campaign targeting the OPRM1 receptor using a modern generative ML pipeline (Appendix A.1, Figure 3a). We diffused 300,000 peptide binder backbones of length 8-10 residues, allocated approximately evenly to ProteinMPNN or PeptideMPNN, and generated a total of 2.4 million sequence designs. Peptide sequences were then filtered after scoring the refolded complexes using AF2. After filtering, approximately 100,000 sequences were assessed for agonism of the OPRM1 receptor with a microfluidics-based functional screening platform from Orbit Discovery. Peptides were subjected to multiple rounds of selection based on receptor agonism activity followed by amplification, and finally sequenced and ranked by frequency. After sequencing, approximately 55% of remaining sequences originated from PeptideMPNN, and approximately 45% originated from ProteinMPNN. Further details are included in Appendix A.1.

We assessed the relative enrichment of the sequence design methods within the top-ranked peptides, as depicted in Figure 2a. PeptideMPNN designs were significantly enriched above base rate occurrence within the peptides exhibiting the highest agonism activity (global *p* = 0.0425 over 10 *<*= *k <* 50 using 2000 random ranking permutations), contributing 9 of the top 10 sequences (p-value= 0.021) and 25 of the top 30 sequences (p-value=0.010). Further details of enrichment factor and p-value calculation are included in Appendices A.2.4 and A.2.5. The AF2 predicted structure of the top-ranked PeptideMPNN design is shown in Figure 2b, exhibiting a favorable binding pose within the orthosteric pocket.

**Figure 2.**
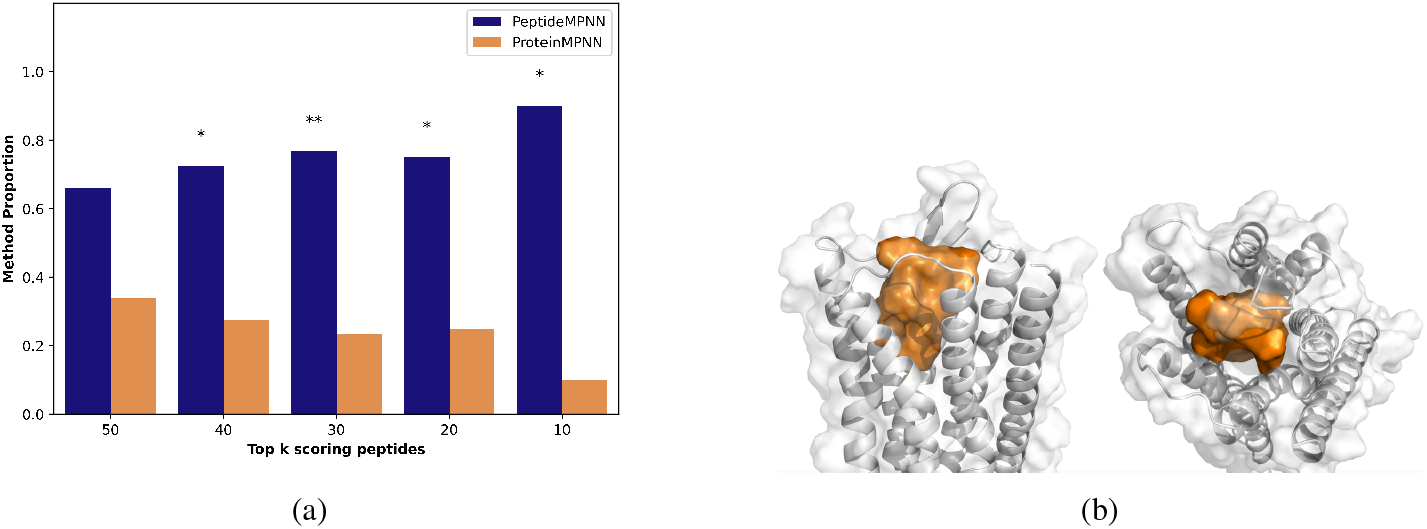
(a) Experimental agonism-selected Next Generation Sequencing (NGS) enrichment within *de novo* peptide designs. Sequences designed by PeptideMPNN exhibit significant enrichment above base rate occurrence (55%) within top ranked peptides, in comparison to ProteinMPNN. Values of *k* marked with “*” or “**” denote enrichment of PeptideMPNN with rank-permutation p-value *<*= 0.05 and *<*= 0.01 respectively. (b) The AF2 predicted structure of the experimentally top-ranked PeptideMPNN design in complex with OPRM1.

**Figure 3.**
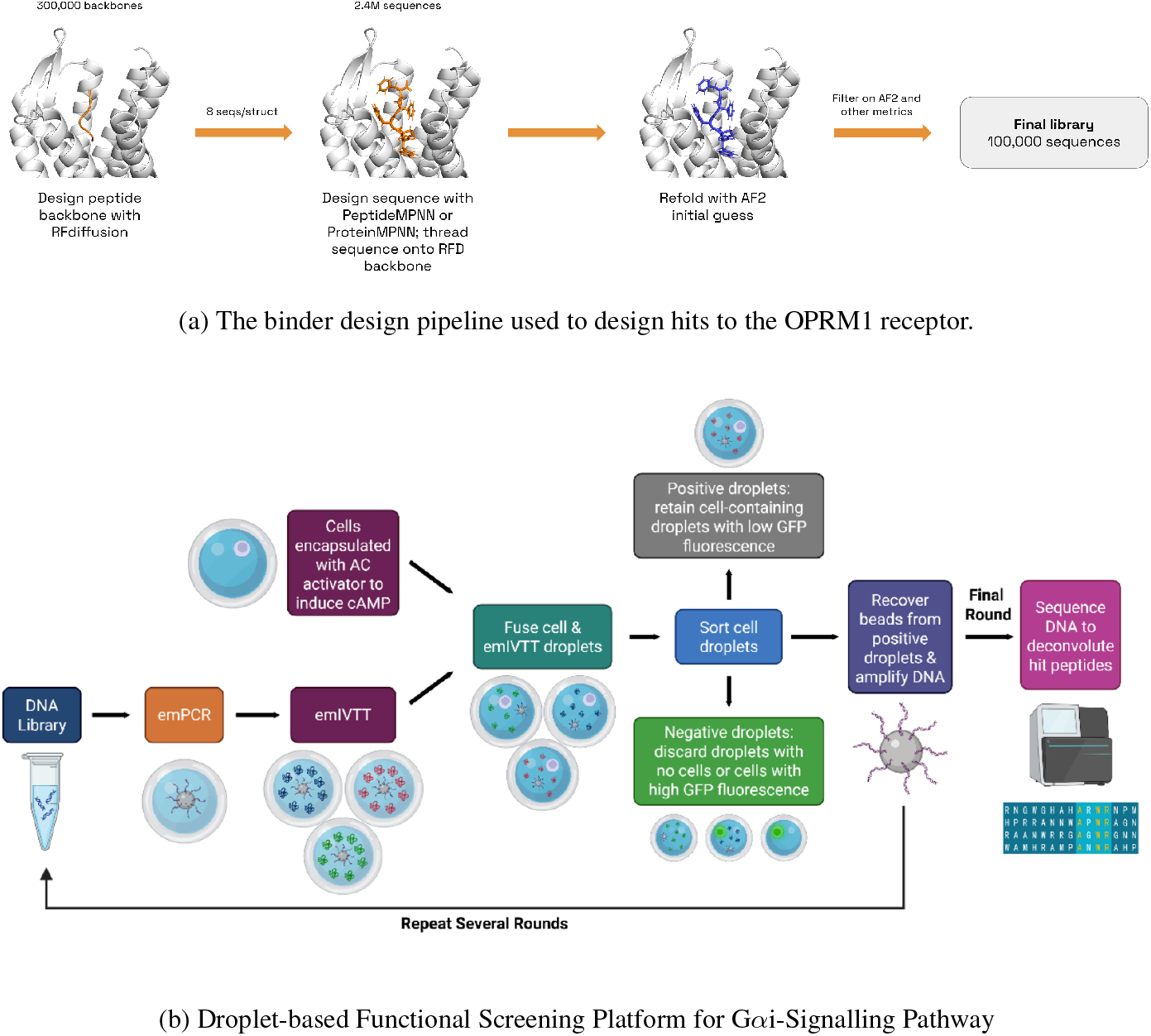
Methods used to design and screen the peptide library against OPRM1.

**Figure 4.**
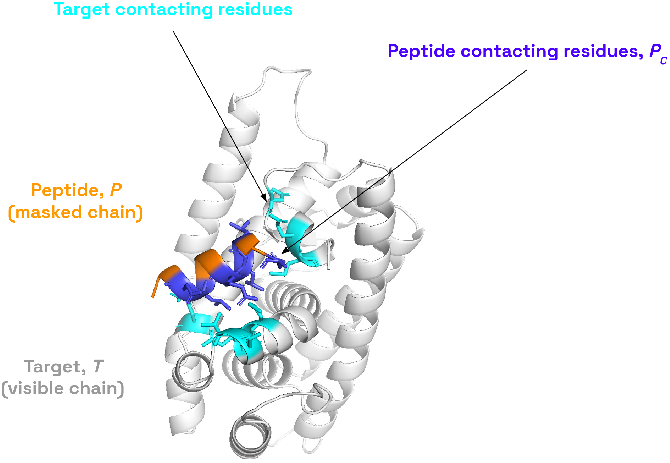
Labeling of contacting residues in a peptide-protein complex.

## Conclusion

Peptides are a growing class of therapeutics. While computational methods are increasingly successful at *de novo* protein design, performance for designing peptide binders lags behind. In this work, we contribute a peptide-specific inverse folding model with the aim of addressing this shortcoming.

We demonstrated that finetuning the inverse folding model ProteinMPNN on peptide-protein complexes results in improved peptide sequence recovery. We find that a training and finetuning regime with elevated levels of backbone noise is beneficial for peptide inverse folding. The benefits of PeptideMPNN were validated *in vitro*, as it produced significantly more peptides exhibiting elevated receptor agonism than ProteinMPNN. Additionally, we present a novel contact-based decoding order for *de novo* peptide design. Contact decoding was found to improve *in silico* success rates for peptide binder designs in comparison to random order decoding by 16% *™* 30%.

Multiple possibilities exist for future work, including exploring different regularization strategies to improve finetuning, considering variations on model parameters such as the number of layers and GNN neighbors, and pursuing additional experimental validation against a broader set of targets.

## A Appendix supplemental material

### A.1 In vitro methods

Our binder design pipeline is summarized in Figure 3a. The experimental assay used to screen for OPRM1 agonism is shown in Figure 3b.

A single codon DNA library, encoding linear 8-10mer peptides, was designed according to the methodologies described in Figure 3b. The library comprised the 8-10 mer peptide encoding region within a flanking scaffold containing all elements required for DNA amplification (via emulsion PCR; emPCR) and translation (via emulsion *in vitro* transcription translation; emIVTT). The DNA-encoded library was procured from Twist and underwent NGS-based QC upon receipt to confirm library quality prior to screening.

CHO-K1 cells reporter cells stably expressing OPRM1, mCherry (as a cell marker) and CRE-GFP (cAMP response element linked green fluorescent protein) were utilised for screening. Application of an activator of adenylyl cyclase (e.g. forskolin or NKH-477) to these cells results in induction of cAMP through the Gs-signalling pathway and eventually production of GFP. Stimulation of OPRM1, which signals via the Gi-signalling pathway, results in inhibition of the Gs-signalling pathway and a reduction in GFP.

Screening was performed using Orbit Discovery’s microfluidics-based functional screening platform 3b. Briefly, the DNA-encoded library is first encapsulated with primer-coated beads and PCR reagent within droplets, to generate a library of DNA-coated beads via emPCR, each presenting a single species of DNA. After QC of the resulting DNA-coated beads, the library beads are subsequently encapsulated with IVTT reagent within emIVTT droplets and incubated overnight. This allows for the production of peptides corresponding to each of the library sequences via IVTT. Peptides are expressed off-bead and remain free in solution within the droplet, maintaining the genotype-phenotype linkage between peptide and the encoding DNA. The resulting peptide-filled droplets are merged with droplets containing cells and NKH-477 (to activate adenylyl cyclase and induce the Gas-signalling pathway via cAMP, resulting in production of a fluorescent GFP signal). The resulting cell and peptide containing droplets are then incubated overnight to allow for peptide-receptor interaction. If a peptide stimulates the receptor, this will result in inhibition of cAMP and a decrease in GFP and the level of fluorescence. Following incubation, droplets are sorted using a droplet sorter based upon two parameters; 1) high mCherry fluorescence – indicating that the droplet contains a cell and 2) low GFP fluorescence – indicating inhibition of stimulation via the Gi-signalling pathway. The DNA-coated beads are recovered from positive sorted droplets and the DNA amplified off bead and assembled back into the full-length Orbit scaffold via overlap extension PCR. This results in a library of enriched DNA, which is then used as the template for emPCR in the subsequent round of screening. Four rounds of screening were performed in this way.

After the final round of screening, purified DNA recovered from each round was prepared for next-generation sequencing (NGS) to deconvolute the identity of hit peptides. The purified DNA underwent a two-step PCR process using custom primers to add the adaptor sequences and indices required for sequencing. The resulting DNA was sequenced with a custom primer using a Miseq Reagent Kit v3 and Illumina Miseq NGS System. The resulting FASTQ files were analysed via a custom pipeline to determine the identity of enriched peptides. Peptides were ranked both by overall frequency after Round 4 selection and by their level of enrichment from Round 1 emPCR to after Round 4 selection.

### A.2 Contact-based decoding order

Figure 4 illustrates how fixed, masked contacting and masked non-contacting residues can be labeled on an input structure.

#### A.2.1 Selection of contact distance threshold, *d*_*c*_

We examined two different expert-suggested C-*α* distance thresholds, *d*_*c*_ = [6, 8]*Å*in our peptide test set. We expected a reasonable contact distance should yield close to zero density when the proportion is zero. Figure 5 shows that at 6*Å*, we observe a subset of peptide-protein complexes which have zero contacts. However at 8*Å*, we see that only one test structure exhibits zero contacts at this threshold. Therefore, we used *d*_*c*_ = 8*Å*.

**Figure 5.**
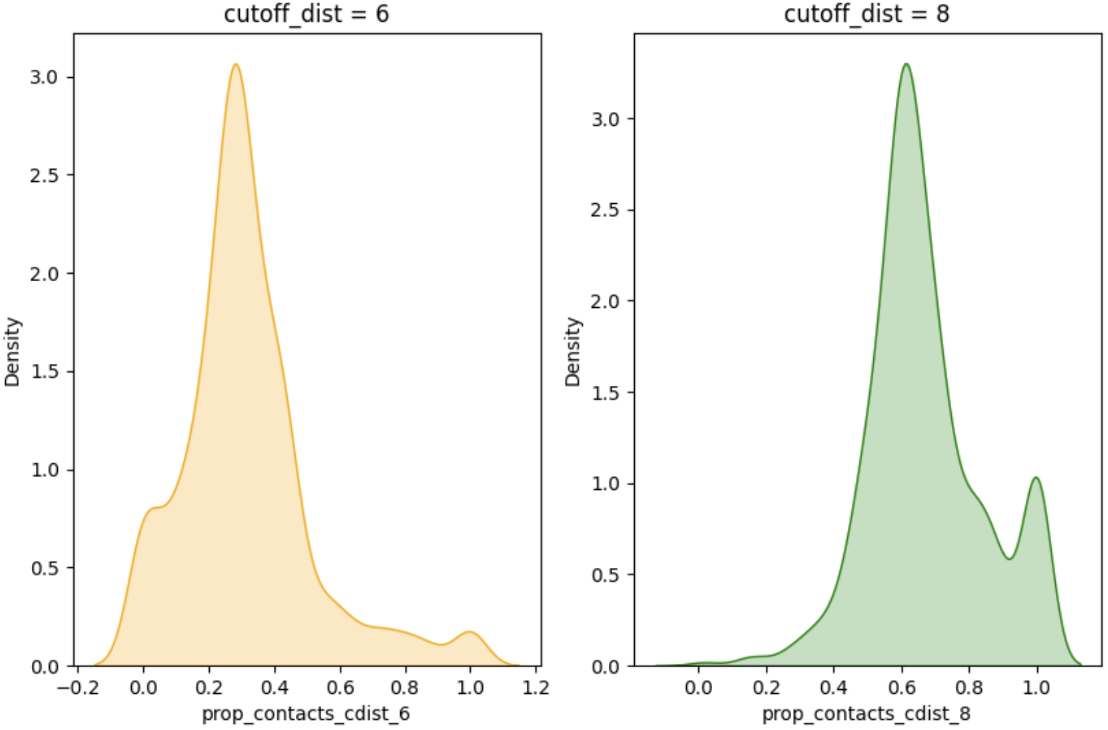
Distribution of inter-chain contacts at different C-*α* distance thresholds. The x axes represent proportions of inter-chain contacts.

**Figure 6.**
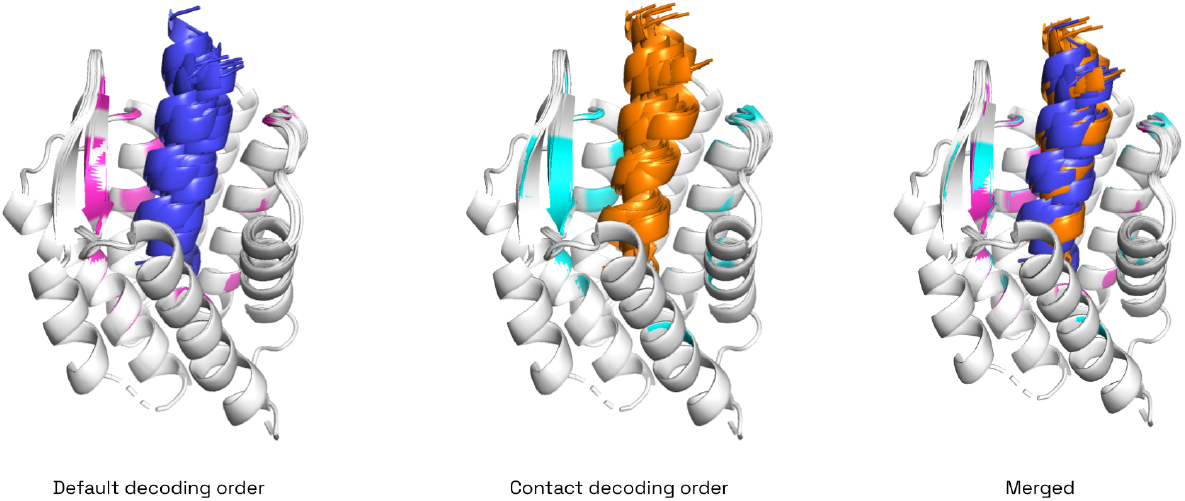
Structures of top-10 *in silico* designs (`pae_interaction` < 10) using the random (left) and contact-based (middle) decoding orders. Right shows random and contact-based designs overlayed. The contact residues on the target (side-chains within 4*Å*) for random and contact-based decoding are coloured in magenta and cyan, respectively.

**Figure 7.**
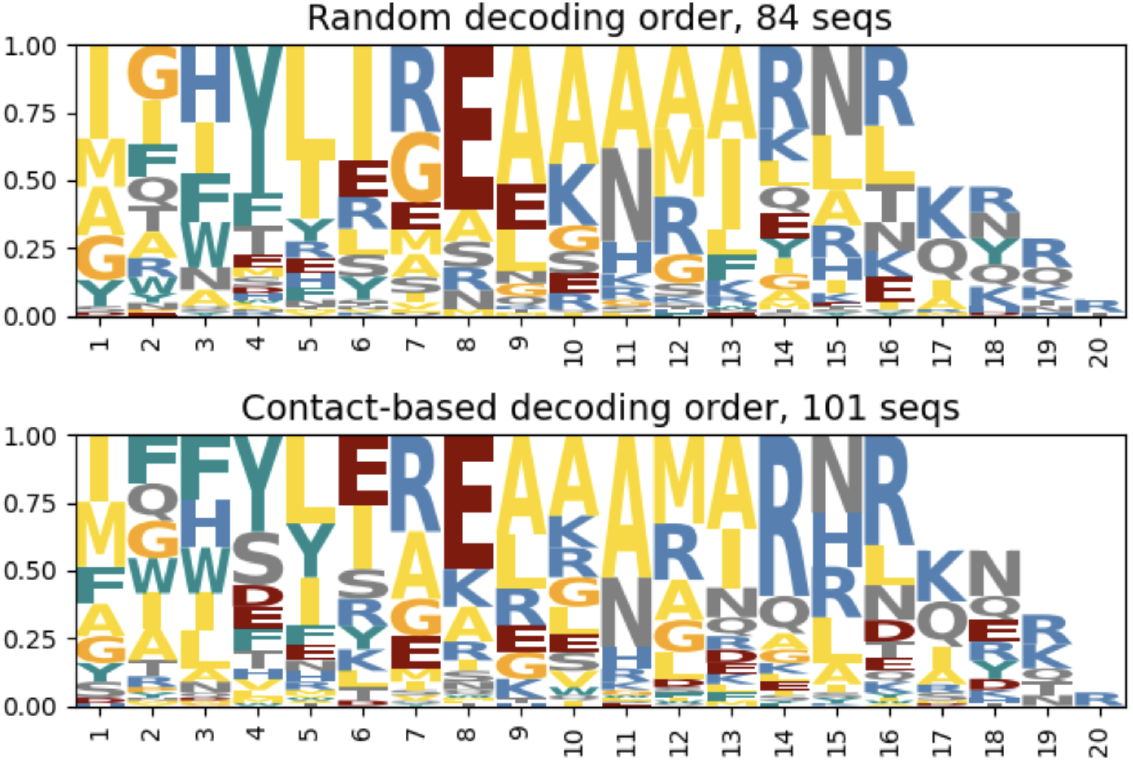
Sequence logos of top-ranked *in silico* designs (`pae_interaction` < 10) using the random (top) and contact-based (bottom) decoding orders.

#### A.2.2 Decoding order performance, complete tables

The full table of test loss and accuracy is shown here in Tables 3 and 4, comparing performance across decoding order strategies on PeptideMPNN models finetuned with the alternative learning rate schedule described in Appendix A.3. Contact-decoding performs almost identically to random decoding.

#### A.2.3 Decoding order effects on *de novo* binder design

The structures of the ten lowest `pae_interaction` designs for each decoding order strategy are shown in Figure 6. Both methods generate helical peptides that bind deep in the OPRM1 pocket. The collective contact-based designs qualitatively appear to make more consistent contact across the surface of the receptor’s *β*-sheet compared to the random decoding order designs.

We N-terminally aligned the *in silico* hits (`pae_interaction` < 10) from *de novo* binder design to the target OPRM1 and generated sequence logos (Figure 7) since we observed that most designs docked the N-terminus in the binding pocket.

Endomorphin (YPWFX) and *β*-endomorphin (YGGFMTSEKSQTPLVTLFKNAIIKNAYKKGE), the native binders of OPRM1, share a YXXF motif at the N-terminus. This motif is also shared with synthetic peptide binder DAMGO (YAGFX). Both random and contact-based decoding do produce tyrosine at position 1 and phenylalanine at position 4, though these are not the dominant amino acids at those positions. Both methods prefer hydrophobic tyrosine at position 4, which could be considered similar to phenylalanine. Interestingly, contact-based decoding uniquely suggests a phenylalanine at position 1, which we do not see in random decoding designs.

#### A.2.4 Enrichment analysis

Enrichment was calculated as over-prevalence of a sequence design method, compared to expectation from random chance, within the top sequenced hits. We tested multiple rank thresholds for a hit to be considered top-ranked. Specifically, we tested enrichment of the two methods within the “top *k*” peptides for all *k* between 10 and 50, inclusive.

A higher Enrichment Factor indicates a method produced more top-ranking receptor agonists. It is calculated as

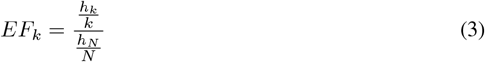

Where *h*_*k*_ is the number of hits from that method in the top-*k* designs, *h*_*N*_ is the total number of occurrences in the full dataset, and *N* is the total size of the dataset. Intuitively, Enrichment Factor at a given *k* is the frequency of a method resulting in the most significant NGS hits, *relative* to its total frequency.

The associated statistical test for Enrichment Factor is the hypergeometric distribution, specifically what is termed the “survival function” or right-tail area *P* [*X > x*]. We parameterized the hypergeom.sf(x, N, K, n) with *x* = *h*_*k*_ − 1, *N* = *N, K* = *h*_*N*_, and *n* = *k*, in reference to the Enrichment Factor equation above. This described the probability of observing at least *h*_*k*_ hits within the top *k* rank by chance, within the context of a single statistical test.

#### A.2.5 Permutation-based multi-test correction

To account for testing for enrichment at various top-*k* thresholds, we executed a permutation-based global null test. The true NGS ranking was randomly scrambled 1999 times, and for each pseudo-ranking the identical sliding-threshold enrichment analysis was performed as described above – fully for all *k* from 10 to 50, inclusive. The number of pseudo-rankings that resulted in at least one test with p-value at least as significant as the minimum true p-value observed were tallied, and used to estimate the probability of achieving this level of Enrichment Analysis (marginal) significance result by random chance.

The final equation for permutation-based global p-value was

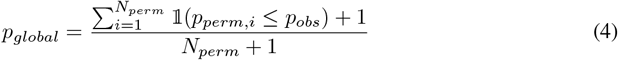

Where *p*_*perm,i*_ is the p-value of the *ith* permutation, *p*_*obs*_ is the p-value observed in the experiment, and *N*_*perm*_ is the total number of permutations performed. One can consider this global p-value as the probability of the null hypothesis *H*_0_ that PeptideMPNN is statistically indistinguishable from ProteinMPNN with respect to likelihood of designing top-ranking sequences. We observed a global p-value of 0.0425.

### A.3 Alternative learning rate schedule

Early experimentation utilized the “NoamOpt” learning rate schedule defined by Vaswani et al. [2017] for finetuning. We found that this finetuning regime was similarly beneficial to finetuning with AdamW for improving sequence recovery, and improved perplexity for models that were both pretrained and finetuned with high levels of backbone noise. However, this LR schedule did not improve on the perplexity exhibited by the best base models. Full results are included in Tables 5 and 6.

**Table 5:** Peptide sequence recovery across pretraining and finetuning *ϵ* values.

| Pretraining $\epsilon$ | No Finetune | FT $\epsilon = 0.02$ | FT $\epsilon = 0.1$ | FT $\epsilon = 0.2$ | FT $\epsilon = 0.3$ |
| --- | --- | --- | --- | --- | --- |
| 0.02 | $0.554 \pm 0.001$ | $0.517 \pm 0.014$ | $0.547 \pm 0.010$ | $0.564 \pm 0.017$ | **$0.559 \pm 0.012$** |
| 0.1 | $0.541 \pm 0.001$ | $0.535 \pm 0.015$ | $0.566 \pm 0.016$ | **$0.572 \pm 0.014$** | $0.569 \pm 0.012$ |
| 0.2 | $0.511 \pm 0.001$ | $0.531 \pm 0.010$ | $0.565 \pm 0.014$ | **$0.590 \pm 0.010$** | $0.583 \pm 0.020$ |
| 0.3 | $0.464 \pm 0.001$ | $0.520 \pm 0.022$ | $0.552 \pm 0.012$ | $0.577 \pm 0.013$ | **$0.587 \pm 0.010$** |

**Table 6:**
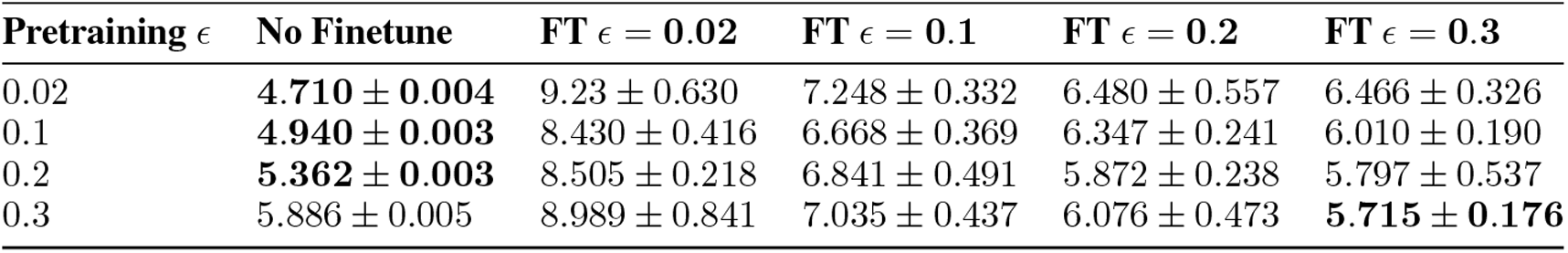
Peptide sequence perplexity across pretraining and finetuning *ϵ* values.

| Pretraining $\epsilon$ | No Finetune | FT $\epsilon = 0.02$ | FT $\epsilon = 0.1$ | FT $\epsilon = 0.2$ | FT $\epsilon = 0.3$ |
| --- | --- | --- | --- | --- | --- |
| 0.02 | **$4.710 \pm 0.004$** | $9.23 \pm 0.630$ | $7.248 \pm 0.332$ | $6.480 \pm 0.557$ | $6.466 \pm 0.326$ |
| 0.1 | **$4.940 \pm 0.003$** | $8.430 \pm 0.416$ | $6.668 \pm 0.369$ | $6.347 \pm 0.241$ | $6.010 \pm 0.190$ |
| 0.2 | **$5.362 \pm 0.003$** | $8.505 \pm 0.218$ | $6.841 \pm 0.491$ | $5.872 \pm 0.238$ | $5.797 \pm 0.537$ |
| 0.3 | $5.886 \pm 0.005$ | $8.989 \pm 0.841$ | $7.035 \pm 0.437$ | $6.076 \pm 0.473$ | **$5.715 \pm 0.176$** |

To further interrogate why the initial finetuned model with both pretraining and finetuning *ϵ* = 0.3 exhibited improved sequence recovery but higher perplexity in comparison to the base model with *ϵ* = 0.02, we compared the distribution of residue probabilities between models, finding that the PeptideMPNN models exhibit more confident, low-entropy predictions (see Figures 8 and 9). Despite this, the increased accuracy was found to be beneficial for *de novo* sequence design tasks, where in practice the entropy levels of class probabilities are controlled by temperature scaling during sampling.

**Figure 8.**
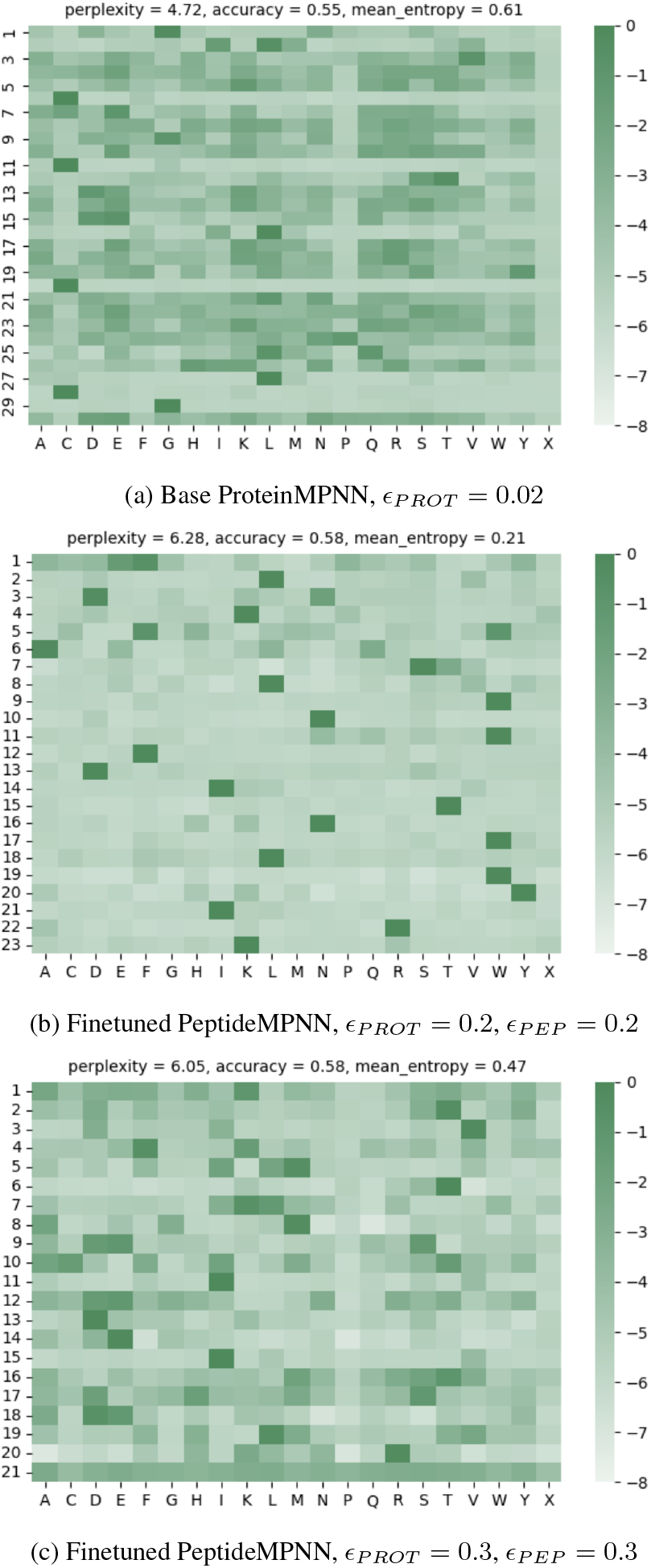
Log probability matrices of the highest perplexity samples from different models finetuned with the initial Noam LR schedule. We observed that the finetuned models tended to be more confident in their predictions, regardless of whether the preferred amino acid was “correct” (native) or “incorrect”.

**Figure 9.**
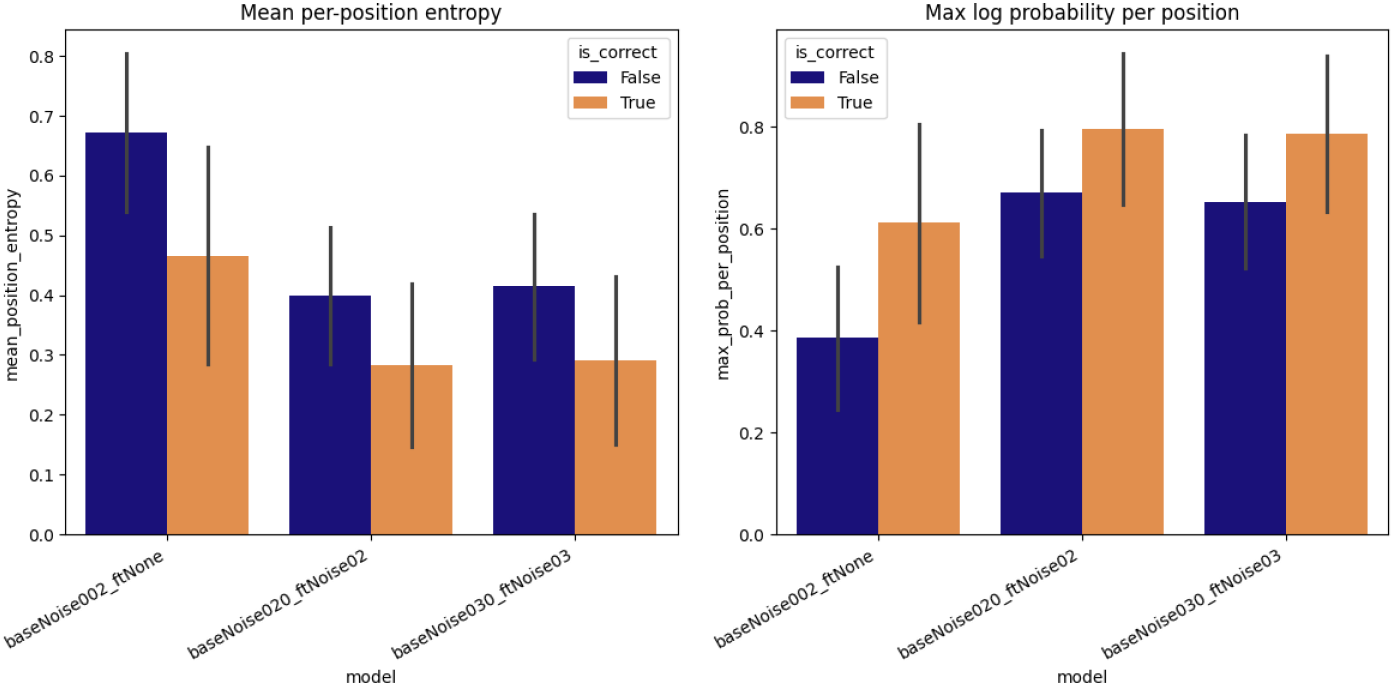
Mean entropy and max probability per position in log probability matrices. *is*_*correct* = *True* indicates the native amino acid had the highest probability, while *is*_*correct* = *False* refers to a non-native amino acid having the highest probability.

This trend was confirmed in aggregate over the test set, shown in Figure 9.

### A.4 Refolding the test set with AF2

In order to assess designability and robustness to imperfect atomic coordinates, Dauparas et al. [2022] tested sequence design on structures generated with Alphafold2 (AF2). They ran all five AF2 ptm models, using only single sequence as input (no templates or MSA), and selected the model with the highest average pLDDT for each input.

We attempted to replicate their AF2 test set generation with AF2 initial guess (AF2IG), which uses a structure of the target with the peptide sequence threaded onto the designed or native peptide backbone as a template. AF2IG does not use MSA. The structure of the target is fixed, while only the structure of the binder is predicted by the model. Because AF2IG uses only one set of model weights, only one structure is produced per input.

We chose to select the highest quality structures by `pae_interaction` rather than pLDDT as done by Dauparas et al. [2022], as the total average pLDDT is largely influenced by the fixed target structure. pae_interaction better reflects the accuracy of the structure at the binding interface. Only 269 structures with `pae_interaction` < 15 were selected to be part of our peptide AF2 test set.

Given that AF2IG uses a template, where Dauparas et al. [2022] used the original AF2 multimer without any templates or MSA, we theorized that AF2IG would produce more accurate structures than what was generated in the original ProteinMPNN paper. To verify this, we looked at RMSD of the true and predicted peptide structures, first aligning by the target (Figure 10 left and middle) to assess prediction of the peptide structure and binding location, and also aligning the peptide (Figure 10 right) to evaluate prediction of the internal peptide structure.

**Figure 10.**
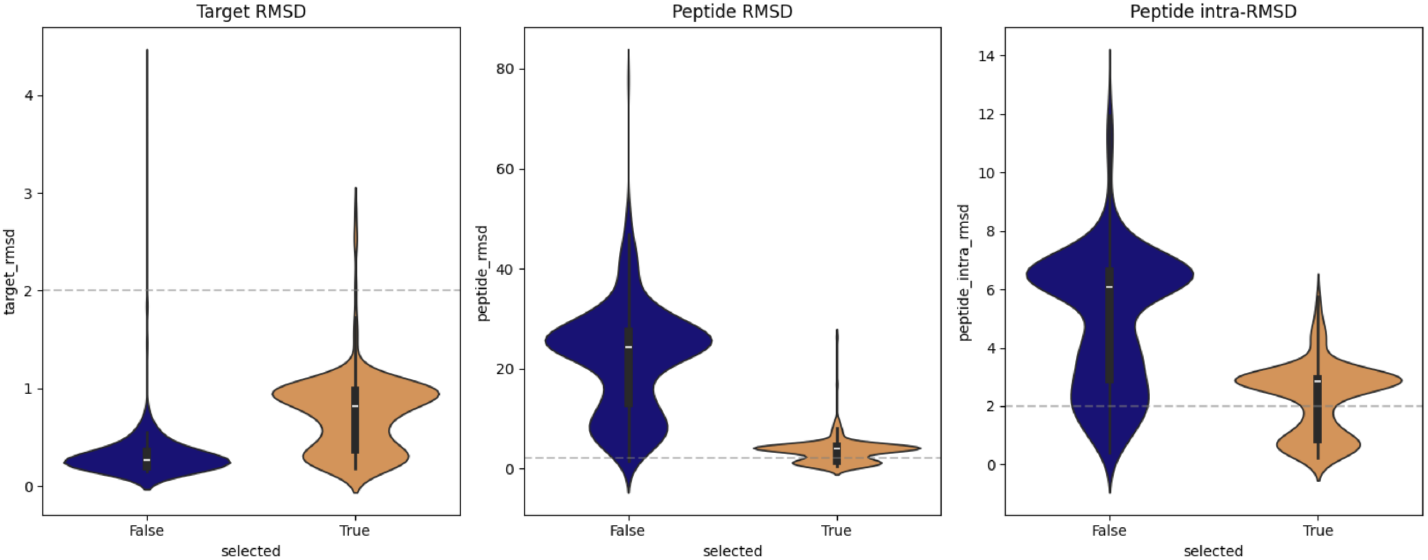
Accuracy of AF2IG on the peptide test set. *selected* = *True* indicates structures which had a `pae_interaction` < 15, of which there were 269 in total. The grey line denotes the maximum acceptable RMSD threshold for what is considered a reasonable prediction. Peptide and target RMSDs are calculated aligning by the target, while peptide intra-RMSD is calculated aligning the peptide by itself.

We observed many predicted structures where the target-aligned peptide RMSD and intra-peptide RMSD were above 2*Å*. The number of high-RMSD structures was lower in the selected set compared to the non-selected set, however the selected set still had a large number of structures with high peptide and intra-peptide RMSDs. This indicates that AF2IG is incorrectly docking the peptide to the target, as well as making errors in the peptide structure itself. Figure 11 shows one such example.

**Figure 11.**
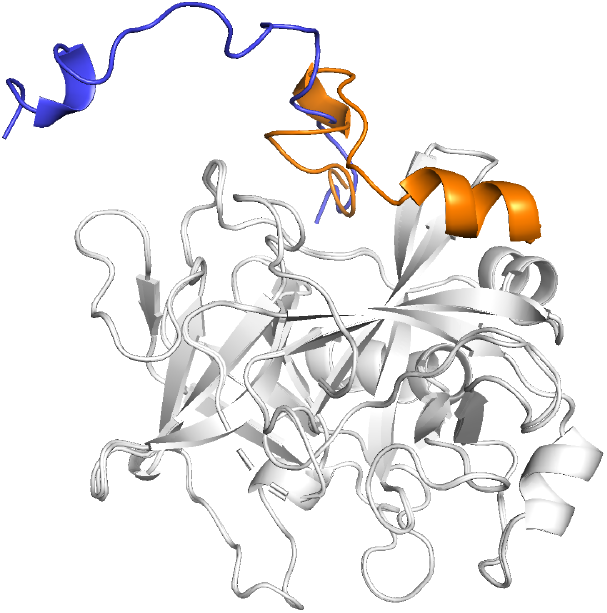
Example of an AF2IG prediction with high target-aligned RMSD (23*Å*) and peptidealigned peptide RMSD (7*Å*). Both the placement of the predicted peptide (blue), and the peptide structure, differ significantly from the true structure (orange).

It is unclear if Dauparas et al. [2022]’s AF2 test set suffered from similar problems, since RMSD to the true experimental structures is not reported. However, given that we are trying to predict peptide structures, which are notoriously more disordered than proteins, it is possible that the inaccurate test set generation is specific to our case.

Based on this, we opted not to use our peptide AF2 test set for benchmarking.

